# Chimeric chikungunya virus-like particles with surface exposed SARS-CoV-2 RBD elicits potent immunogenic responses in mice

**DOI:** 10.1101/2023.01.29.526074

**Authors:** Vedita Anand Singh, Sanket Nehul, Chandra Shekhar Kumar, Manidipa Banerjee, Pravindra Kumar, Gaurav Sharma, Shailly Tomar

**Affiliations:** Department of Biosciences and Bioengineering, Indian Institute of Technology Roorkee, Roorkee-247667, Uttarakhand, India; Kusuma School of Biological Sciences, Indian Institute of Technology – Delhi, Hauz Khas, New Delhi 110016, India; Indian Veterinary Research Institute, Izatnagar, Bareilly, Uttar Pradesh state - 243122, India

**Keywords:** Chimeric enveloped virus-like particles (Chi-eVLPs), chikungunya virus (CHIKV), SARS-CoV-2, Receptor Binding Domain (RBD), Envelope glycoprotein, hybrid VLP vaccine

## Abstract

The SARS-CoV-2 pandemic has reinforced efforts for developing effective vaccination strategy for existing and emerging viruses. Currently there are various vaccine technology available for treating viral diseases, however it is imperative to develop and investigate second-generation vaccines such as chimeric virus-like particles (chi-VLPs) vaccine for increased immunogenicity, ease of production and scalability to supplement the worldwide vaccine supply. Intriguingly, chi-VLPs expresses more than one antigenic epitope on its surface, hence it is expected to be a more effective vaccine candidate. Hereby, this study reports, a novel bivalent vaccine design of chimeric alphavirus coronavirus virus-like particles (ChAC-VLPs), displaying fusion glycoproteins of CHIKV and receptor binding domain (RBD) of SARS-CoV-2 on its surface. Uniqueness and versatility of ChAC-VLPs has been demonstrated via a various techniques including Western blot, Immunofluorescence, cryoEM, and dynamic light scattering (DLS). The multimeric epitope display of immunogenic antigens, i.e CHIKV envelop glycoprotein and SARS-CoV-2 RBD was validated by cell-based assays. ChAC-VLP immunized mice has shown substantial neutralization titres for CHIKV (PRNT50 of 1:25) from the serum collected after 2^nd^ booster doses. Similarly, serum antibodies were detected for SARS-CoV2 RBD as observed by antigen specific ELISA and validated using surface plasmon resonance (SPR). SPR binding response was detected to be >200 RU for anti-RBD antibody in post-immunized mice sera. In conclusion, present study proposes ChAC-VLPs as a potential hybrid vaccine candidate for CHIKV and SARS-CoV-2 infection and contributes valuable insights in chi-VLPs domain.

## 1. Introduction

Since the advent of VLPs (Virus-like particles), a wide variety of VLP-based vaccines have been developed for the prophylactic treatment of numerous infectious agents (1). After the approval and commercialization of VLP-based vaccines, Engerix-B® (Hepatitis B Virus) and GARDASIL®9 (Human Papillomavirus Virus, Quadrivalent), a numerous VLP candidates have been studied since then, some of which have entered clinical trials (1, 2). The versatile VLP technology introduces dense repetitive array of structural proteins of wild type viruses. These units remarkably resemble wild type viruses without being infectious. The use of eukaryotic expression system has allowed researchers to develop various prophylactic VLP-based vaccine candidates for a few highly pathogenic viruses such as HIV (3), severe acute respiratory syndrome (SARS) (4–6), Ebola virus (7, 8), dengue fever virus (9–11), chikungunya virus (CHIKV) (12, 13), Rift Valley fever virus (RVFV) (14) and HCV (15, 16). Vaccines that use VLPs are safer than inactivated or attenuated viruses as the chances of virus reversion or incomplete inactivation is avoided. VLPs based vaccines are known to improve the effectiveness of immunological stimulation by increasing the active components of the immune response hence stimulating the body’s natural defences (17–20). Moreover, VLPs have the advantage of being more effective than other subunit vaccines due to their enhanced immunogenic responses (21, 22). Interestingly, VLPs can be produced in various configurations which renders them easily recognizable by APCs (Antigen presenting cells) (22). VLPs can also be produced in a wide range of expression systems including bacterial, yeast and mammalian expression systems which provides flexibility in modifying manufacturing conditions (23, 24). The development of a commercially sustainable and stable VLP vaccine depends on optimising the pH, ionic strength, salt composition etc of the storage buffer, which is crucial for maintaining the integrity of VLPs in solution and its export and import into different countries (25, 26). Structurally, VLPs have been classified into three major classes based on their structural diversity: enveloped, non-enveloped, and chimeric (27).

A chimeric VLP (chi-VLP) is generated by adding antigenic epitopes of more than one virus in a carrier vector hence triggering multivalent immunostimulatory responses (28). The use of chimeric virus-like particles (chi-VLPs) as vaccines and gene vectors has become increasingly significant in both fundamental and applied research. It is intriguing that this technique would offer a platform for a vaccination that could target several pathogenic viruses in a single vaccine formulation. Recent clinical studies have demonstrated the effectiveness of an immunologically active recombinant MeV-based CHIKV vaccine for the treatment of healthy adult patients without a known history of anti-meV immunity (29). Due to their capacity to elicit cytolytic T lymphocyte (CTL) responses, chi-VLPs have gained significance in the development of next-generation vaccines (30, 31). VLPs can be categorised structurally into enveloped (eVLPs) and non-enveloped (non-eVLPs) types (23). Non-eVLPs such as HPV VLPs, has a comparatively simple structure that is made up of one or more structural proteins of the native virus (2, 32, 33). eVLPs have a far more complicated composition than non-enveloped VLPs due to the presence of host cell derived membranes which provides more opportunities for the integration of heterologous antigens and adjuvants (2, 34, 35). The present study demonstrates the design, production and mice immunization studies on chi-eVLPs of CHIKV structural proteins with antigenic epitope comprising of receptor binding domain (RBD) of SARS-CoV-2 and CHIKV E2 glycoprotein.

CHIKV belongs to genus alphavirus genus which comprises of enveloped, single stranded, positive sense RNA genome (36–38). The post-chikungunya status shows similarities with the long-term effects of the COVID-19 infection in terms of socio-economic effects and overall disease burden (39). Currently, licenced no antiviral therapy or vaccine is available for any alphavirus infection worldwide (40–42). Hence, the only effective way to control the spread of these viral infection is through vaccinations in endemic areas and the most vulnerable populations. However, the use of attenuated virus vaccines has its limitations and safety issues. During the numerous waves of the COVID-19 pandemic, the protection provided by existing vaccinations has dramatically diminished because of the rapid emergence of various SARS-CoV-2 variants (43, 44). These SARS-CoV-2 subtypes continue to pose a risk to public health across the world. Therefore, it is essential to establish more effective vaccines and immunization techniques to manage any unpredictable resurgence of SARS-CoV-2 variants and re-emerging CHIKV infections. The emerging lineages of COVID-19 of which SARS-CoV-2 B.1.1.529 (Omicron) variant has 15 mutations in the receptor-binding domain (45). The ChAC-VLP (chimeric alphavirus coronavirus virus-like particles) plasmid design opens avenues to generate stable mature immunogenic VLP platform. The ChAC-VLP plasmid design will give us flexibility to replace and add mutated RBD of any new variant of COVID-19. The goal of our invention is design and develop immunogenic chi-eVLP composition which will elicit immunity against CHIKV and SARS-CoV-2 and its future variants. The ChAC-VLP design enables insertion of the immunogenic domain of any pathogenic viral or bacterial protein, to be expressed as fusion protein with envelop glycoprotein of alphaviral E2. This will pave the way for hybrid VLP vaccine technology wherein resulting single VLP vaccine will protect individuals against more than one pathogenic organism.

## 2. Results

### 2.1. Expression and purification of ChAC-VLPs from HEK293Ts

The structural gene cassette (C-E3-E2-6K-E1) of CHIKV 181/25 was cloned into pcDNA3.1 vector. To generate 1:1 fusion protein of RBD-E2, RBD was inserted after the furin cleavge site before the n-terminus of E2 (Figure 1A). Even though RBD is a large insertion in the E2 protein of CHIK, we observed successful packaging of RBD-E2 as a component of ChAC-VLPs. SDS-PAGE, western blot analysis (Figure 1C) and indirect immunofluorescence assay (IFA) were performed to validate the expression of structural proteins of ChAC-VLP (Figure 1B). IFA results demonstrate the of co-expression of RBD and E2 proteins in transiently transfected HEK293T cells using anti-RBD and anti-alphavirus antibody 24 h after transfection. As compared to the wild type CHIKV 181/25, the presence of a protein band at ∼75 kDa indicates expression of a fusion protein of RBD and E2, which would have otherwise been observed at ∼55 kDa for E2. This was further validated by western blot using anti-RBD antibody. Cryo-electron microscopy revealed a spikey spherical morphology for ChAC-VLPs. Dynamic light scattering (DLS) analysis indicated that the hydrodynamic diameter of the highest intensity peak is 158.9 nm with a PDI of 1.0 although the presence of larger aggregates was also detected. The aggregate peak 2 had a significantly lower intensity compared to the VLP peak (peak 1).

**Figure 1:**
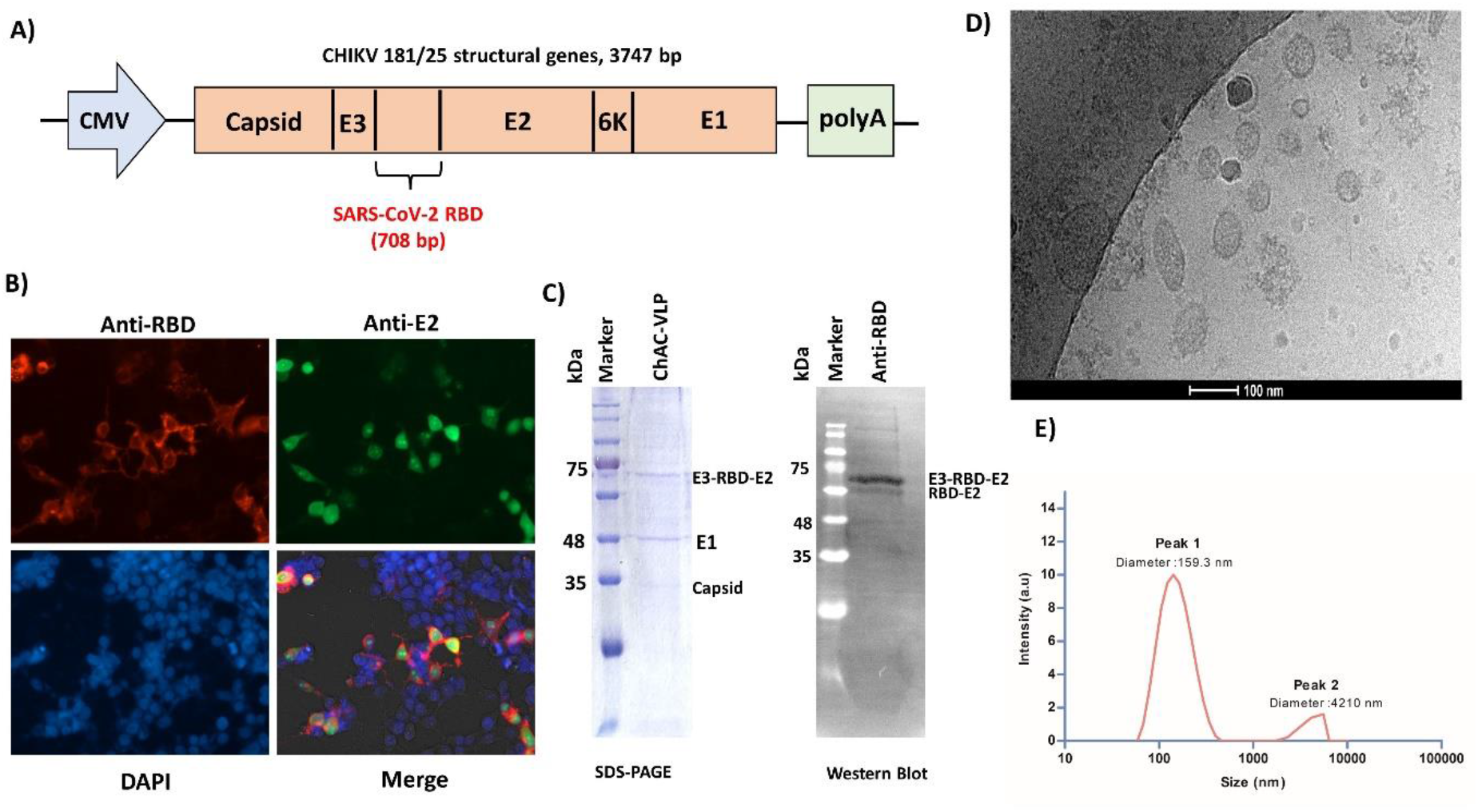
Design, expression and purification of ChAC-VLPs. **A)** Schematic illustration of ChAC-VLP vector design for expression of ChAC-VLP HEK293Ts. The full-length CHIKV 181/25 (vaccine strain) structural gene encoding C-E3-E2-6K-E1 containing SARS-CoV-2 RBD gene was cloned in pcDNA3.1 (+) vector. **B)** Indirect Immunofluorescence images showing the co-expression of RBD and E2 in ChAC-VLP transfected HEK293T cells at 24 hpi. Detection of CHIKV E2 (FITC), SARS-CoV-2 RBD (Alexa 594) and cell nuclei (DAPI) were stained in green, red and blue respectively. Scale bars. ∼50 µm. **C)** To verify the co-expression of CHIKV envelop proteins and SARS-CoV-2 RBD protein in purified ChAC-VLP, SDS-PAGE and Western blot was performed using anti-RBD antibody **D)** Assessment of ChAC-VLP particle size and uniformity using cryo-electron microscopy images of ChAC-VLP particles depicting the presence of purified VLPs. Scale bars. 100 nm. **E)** DLS analysis of ChAC-VLPs with respect to intensity and size of the particles. Peak 1 represents VLP peak and peak 2 represents larger aggregates in purified ChAC-VLP sample.

### 2.2. Multimeric epitope display and antigenic characterization of ChAC-VLPs

#### 2.2.1. Assessment of competitive binding of ChAC-VLPs with CHIKV and SARS-CoV-2 host cell entry receptors

Among the *Alphavirus* genus, CHIKV is clinically the most important member. Like other alphaviruses, Chikungunya virus can infect a wide variety of cells and tissues as the receptor for CHIKV is ubiquitously expressed in host cells. In a previous study, it has been shown that the mCherry insertion at N-terminal of E2 did not cause significant defects in virus assembly and entry (40, 46). Our ChAC-VLP vector design is such the multimeric epitopes (RBD-E2) is exposed on the surface of VLPs. In this work, plaque reduction assay (Figure 2A) and qRT-PCR (Figure 2B) results demonstrates the decrease in CHIKV titres inside host cells in ChAC-VLP treated sample. Similarly, qRT-PCR results show a significant reduction in viral mRNA at 24 hpi in the ChAC-VLP treated cells. Hence, these experiments demonstrate the orientation of RBD-E2 fusion protein on ChAC-VLPs is correct, as they mimic CHIKV surface glycoprotein for entry into host cells as significant reduction in CHIKV viral titre is analysed by plaque reduction assay and qRT-PCR results.

**Figure 2:**
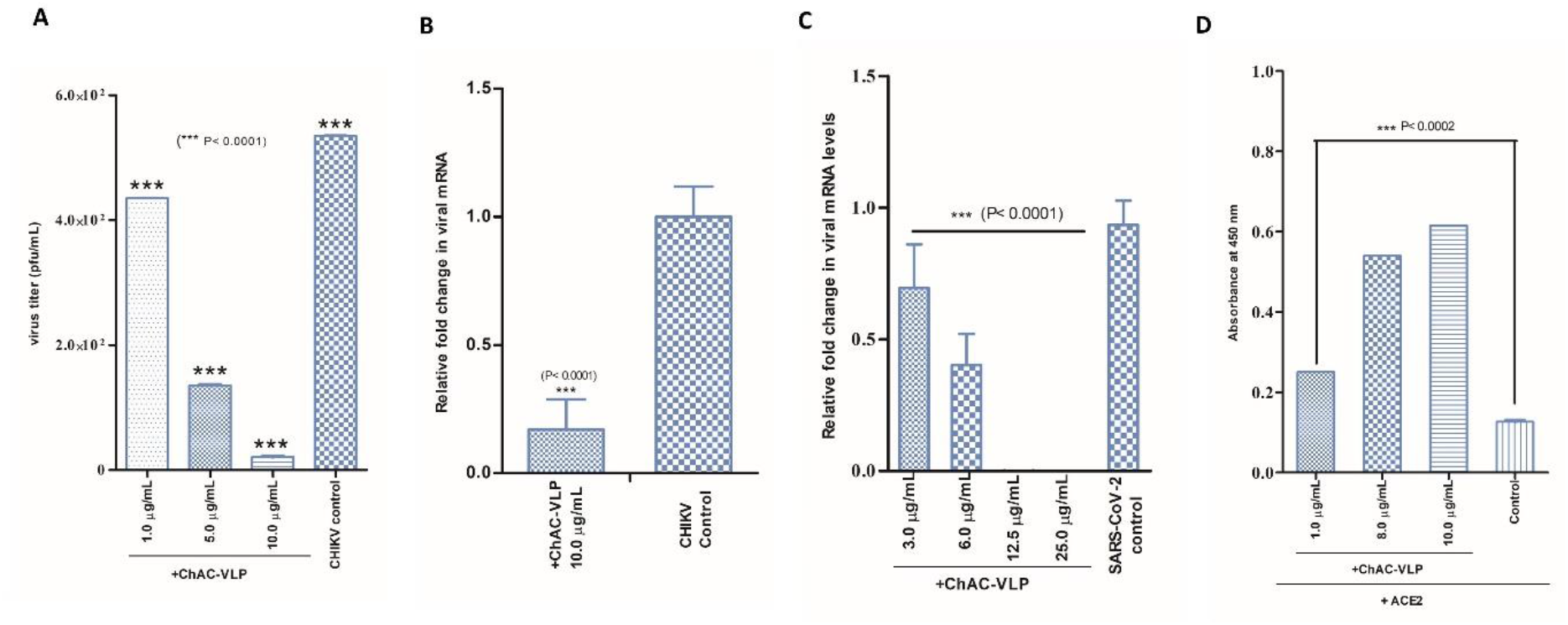
**A)** Reduction in CHIKV titre due to competitive binding of ChAC-VLPs for cell entry receptors shown through plaque reduction assay. Values are mean and error bars are Standard Deviation (n=3). CHIKV plaque images showing the decrease in PFU/mL of ChAC-VLPs treated and non-treated Vero cells and histogram representation of viral titre (PFU/mL) and varying concentration of ChAC-VLPs **B)** RNA was isolated from ChAC-VLPs treated/non-treated cells infected with CHIKV at 24 hpi and reduction in viral mRNA was quantified using RT-PCR. Fold-reduction in viral RNA in ChAC-VLP in treated cells relative to the control is plotted. Actin: endogenous control; CHIKV E1: target (gene. ***, p< 0.001.) Values are mean and error bars are Standard Deviation (n=3). **C)** RNA was isolated from ChAC-VLPs treated/non-treated cells infected with SARS CoV-2 at 48 hpi and reduction in viral mRNA was quantified using one step qRT-PCR. Calculated Ct values were converted to fold-reduction (2^ΔCt) of ChAC-VLPs treated samples compared to non-treated samples. Human RNase P: endogenous negative control; E gene of SARS-CoV-2: target gene. ***, p< 0.001. **D)** ChAC-VLP binding with ACE2 is not affected by RBD incorporation into CHIKV structural proteins. Binding of varying concentration of ChAC-VLP or blank (buffer) to ACE2 in an ELISA format using anti-RBD antibody. Bars represent the absorbance at 450nm mean of duplicates samples with a standard deviation (***, p< 0.001). Values are mean and error bars are standard deviation (n=3).

Furthermore, the SARS-CoV-2 RBD in ChAC-VLP is available as antigenic epitope and competes for the host cell entry receptors with the SARS-CoV-2/human/Ind/CAD1339/2020 virus. Similarly, qRT-PCR results show a significant reduction in SARS-CoV-2/human/Ind/CAD1339/2020 viral mRNA in ChAC-VLP treated cells. This suggests that RBD in RBD-E2 fusion protein is exposed for interaction with ACE2 receptors present on host cell surface. At 48 h, there is >99% reduction in viral RNA present in the supernatant (indicative of released virions) of samples treated with 100 µg of ChAC-VLP compared to control cells (Figure 2C).

#### 2.2.2. ACE2 binding analysis

To validate the correct protein folding of the RBD-E2 fusion protein in purified ChAC-VLPs, ELISA was performed for binding of ChAC-VLP to ACE2 receptor protein. Purified ChAC-VLPs showed efficient binding with the ACE2 which was detected using anti-RBD antibody and HRP-tagged anti-mouse secondary antibody. The goal of the experiment was designed to mimic the SARS-CoV-2 interaction with the host cell in an ELISA format as previously shown by researchers for surrogate virus neutralization test for SARS-CoV-2 (47). The interaction of ACE2 and RBD in ChAC-VLP is observed in dose dependent manner which ensures correct orientation of RBD protein in purified ChAC-VLPs (Figure 2D).

### 2.4. Immunogenicity assessments of ChAC-VLPs in Mice

To evaluate the immunogenicity of ChAC-VLP, Swiss albino mice were immunized with purified ChAC-VLP at Day 0 (prime, 15µg), Day 21 (1^st^ boost, 7µg) and Day 42 (2^nd^ Boost, 5µg). Immunogens were adjuvanted with Alum and injected intramuscularly. The final bleed was performed after 7 days after 2^nd^ boost and sera was collected and processed for immunogenicity assessments. (Figure 3A)

**Figure 3:**
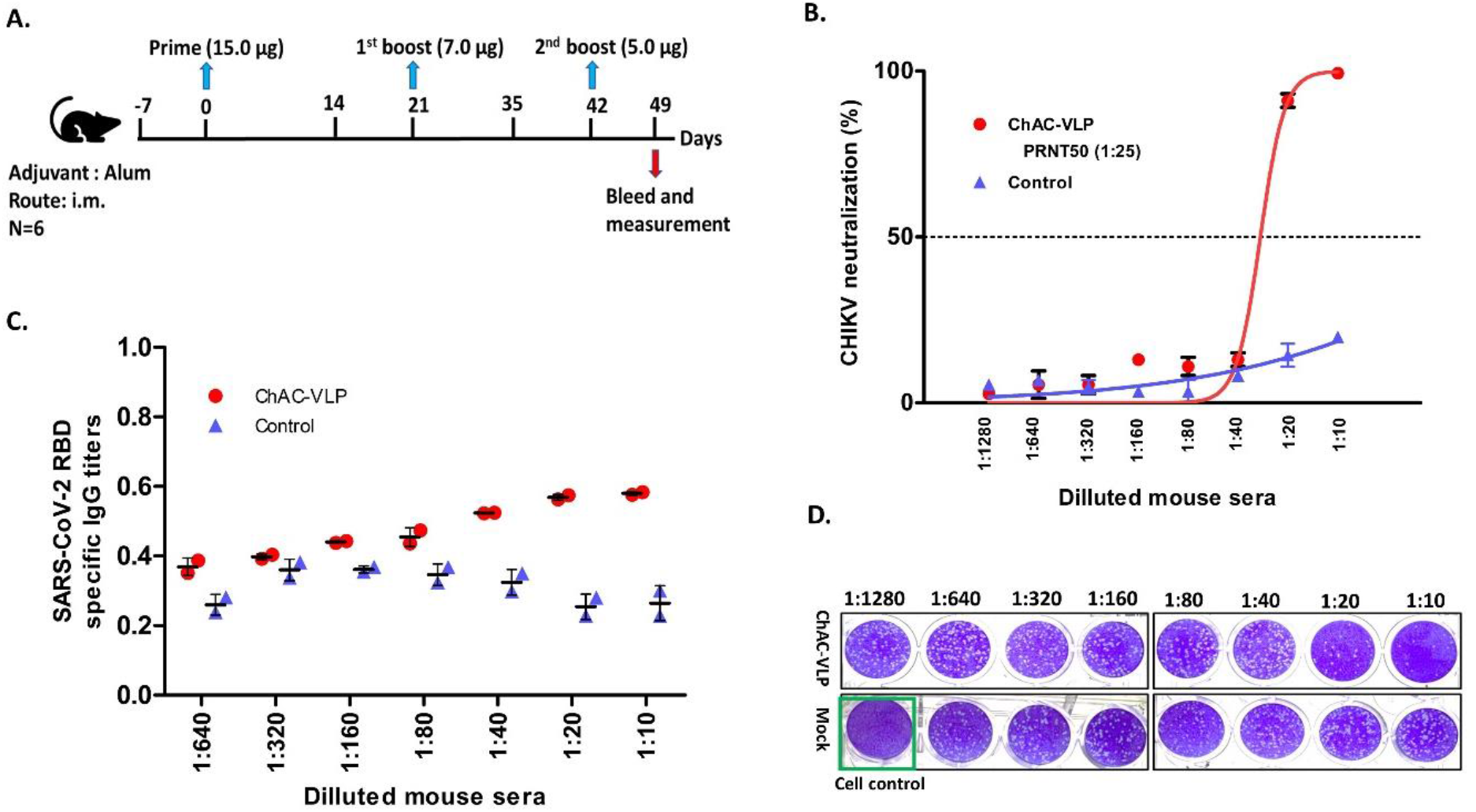
Immunization of ChAC-VLP induced potent immune response against SARS-CoV-2 RBD and CHIKV in mice. **A)** Scheme of the immunization regimen. The control immunogen was alum diluted in PBS and was administered in 2 mice. ChAC-VLP immunogen was administered via i.m route in 4 mice. **B)** Assessment of CHIKV-binding affinity of immunized mice sera: Plaque reduction (PRNT) assay for CHIKV neutralization by anti-CHIKV neutralizing antibody in ChAC-VLP immunized mice sera. The “control” represents the result obtained by infection of virus with addition of control mice sera. CHIKV was tested against serially diluted concentrations (1:10 to 1:1280) of ChAC immunized mice sera and control mice sera. Neutralization dose–response-the PRNT50 data for each antibody dilutions were used to calculate % reduction in plaques. **C)** Detection of anti-RBD antibodies in ChAC-VLPs immunized mice using purified RBD protein coated wells by ELISA. Sera of mice immunized with Alum and PBS were used as control. The data are presented as mean A450 ± SD. (n = 3). The IgG antibody (Ab) titers were calculated as the endpoint dilution that remains positively detectable for SARS-CoV-2 RBD (Red), or control (blue) binding to purified RBD protein. **D)** The PRNT50 values were measured to predict the neutralizing potency of ChAC-VLP immunized mice sera by evaluating the reduction of CHIKV plaques in Vero cells shown as representative plaque images. The results shown are the average±the s.d. of three replicates.

Mouse sera were analysed for SARS-CoV-2 RBD-specific IgG titers by ELISA and neutralizing antibodies for CHIKV was calculated using PRNT50 (Figure 3). Post 2^nd^ Boost, ChAC-VLP elicited sufficiently high RBD-specific IgG titres compared to that of control mice sera. Neutralizing antibody titres were also assessed for CHIKV, and as per our results 1:20 and 1:10 serum dilution showed promising neutralizing activity (Figure 3C). CHIKV PRNT50 was calculated to be 1:25. Even at low doses of ChAC-VLP, the chimeric immune response is generated. The merits of using chimeric VLP as vaccine is the antigen presentation as multiple copies on surface of VLPs as in ChAC-VLP plasmid design, 1:1 incorporation of RBD-E2 is ensured as per previously reported study on reporter-tagged chikungunya virus-like particles (40) and fluorescent protein-tagged Sindbis virus E2 glycoprotein (48). ELISA and SPR was performed to analyse and validate RBD-specific antibody generated in immunized mice sera. For SPR, purified RBD protein was captured on Ni-NTA chip in its native form with a RU value of ∼ 250. The response unit for 2 (Immunized 2 and 4) of the immunized mice was twice as compared to control, whereas the other two mice (Immunised 1 and 3) showed thrice reading compared to the control sera. Thus, the SPR results indicate immunization has induced anti-RBD antibodies in the mice (Figure 4). The results depicted here is immunological assessment of ChAC-VLP in low doses in small mice group. Intriguingly we have observed that ChAC-VLP is efficient in inducing immune response against CHIKV and SARS-CoV-2 RBD.

**Figure 4:**
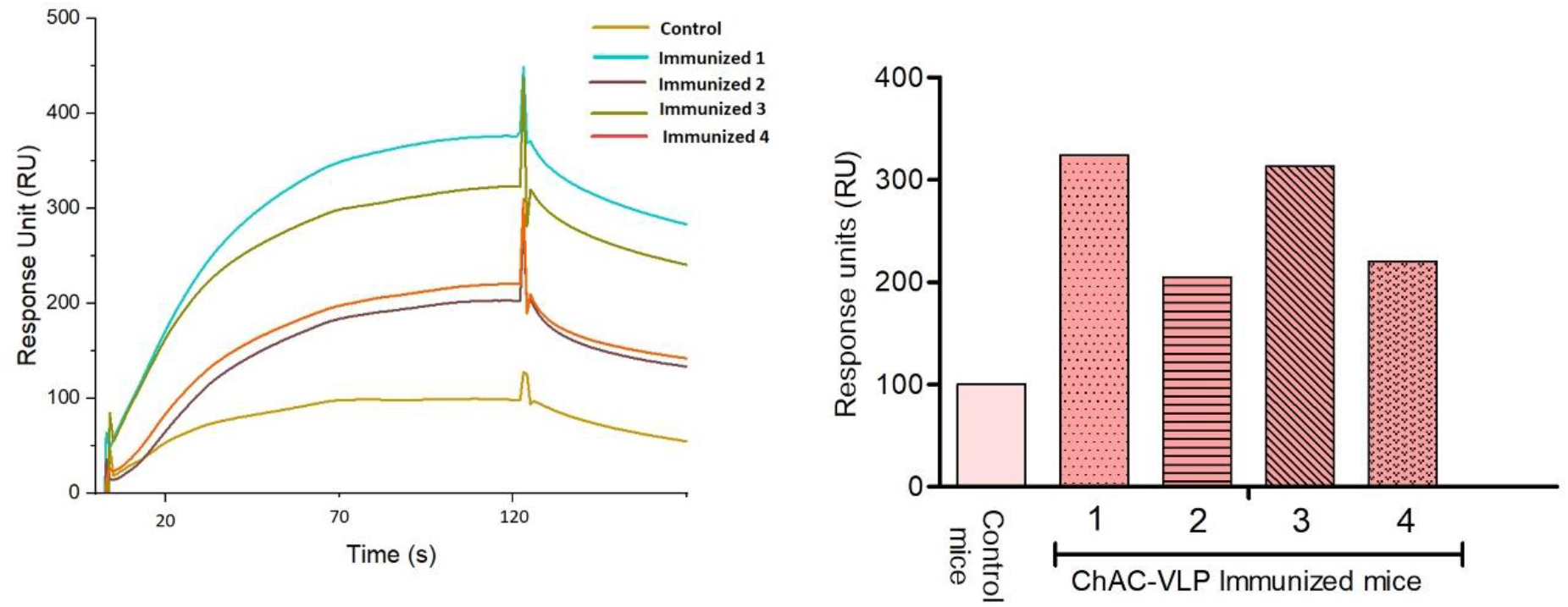
Binding of serum antibodies to recombinant RBD present in immunized mice as compared to non-immunized (Control) mice sera using SPR. The antibody binding is depicted in the form of response unit for 1:10 diluted serum samples. (A) The sensorgram clearly showed the difference between response units of control sera (∼ 100 RU) as compared to the response unit of immunized mice. (B) The sera of immunized mice 1 and 3 showed three times higher readings as compared to the control with the values of ∼ 325 RU and ∼ 315 RU, respectively. Whereas sera of immunized mice 2 and 4 showed two times higher readings with respect to control sera with the values of ∼ 205 RU and ∼ 220 RU, respectively.

## 3. Materials and Methods

### 3.1. Cell line and Viruses

Human embryonic kidney cells (HEK 293Ts, NCCS India) and African green monkey kidney cells (Vero and Vero E6; NCCS India) were grown at 37°C in Dulbecco’s modified Eagle’s medium (DMEM, HiMedia India), supplemented with 10% fetal bovine serum (FBS, Gibco USA) and 1% penicillin-streptomycin (Gibco USA). CHIKV 181/clone25 strain was obtained as a kind gift from Richard J. Kuhn (Purdue University, USA), was propagated and titrated in Vero cells.

### 3.2 Plasmid construction

SARS-CoV-2_S-2P antigen, prefusion spike ectodomain, a gene encoding residue 1-1208 of 2019-nCoV S (GenBank: MN908947) is inserted in n-terminal of E2 protein of CHIKV 181/25 structural genes (C-E3-E2-6K-E1) and cloned into pcDNA3.1(+) expression vector using gibson assembly cloning kit (NEB, USA). The resulting plasmid is designated as ChAC-VLP. A schematic illustration of ChAC-VLP vector design, expression is shown in Figure 1.

### 3.3. Production and Purification of ChAC-VLPs

One day before transfecting HEK293Ts (NCCS, Pune, India), cells were sub-cultured in 1X DMEM supplemented with 10% FBS and 1X penicillin-streptomycin solution in five 150 mm dishes at a seeding density of 5.0 × 10^6^. 2 h prior to transfection, the cell supernatant was removed and fresh 1X DMEM supplemented with 5% FBS and 1X penicillin-streptomycin was added to the dishes. 8 h after transfection, media was changed to fresh 1X DMEM supplemented with 1% penicillin-streptomycin solution. Transfection mix was prepared using calcium-phosphate method and 40 µg of plasmid was added to each 150 mm dish for efficient transfection. VLPs were purified after 72 h post-transfection (hpt). The harvested cell supernatant, was first clarified by centrifugation at 3000 rpm for 30 min, followed by filtration using sterile 0.45 µm PVDF filter (Millipore, USA). For further purification, the clarified supernatant was placed onto a 22% sucrose (Sigma-Aldrich) cushion and centrifuged using SWTi41 rotor (Beckman Coulter Inc., USA) at 110,000*g for 1 h at 4°C. After sucrose cushion, the translucent VLP pellet was gently resuspended in 1X TNE buffer (pH 7.4) (0.1 M Tris-Cl (pH 8.0) 0.01 M EDTA (pH 8.0) 1 M NaCl). ChAC-VLPs was further concentrated to 5 times of its initial volume using 100 kDa cut-off concentrator (Merck, Millipore, USA) followed by analysis on 10% Tris-Glycine SDS-PAGE and successful RBD-E2 fusion protein is validated by western blot using anti-RBD antibody.

### 3.4. Immunofluorescence and Western Blotting

Immunofluorescence assay (IFA), SDS-PAGE and Western blot, and were performed to determine the co-expression of CHIKV structural proteins and SARS-CoV-2 RBD protein. For IFA, ChAC-VLP plasmid transfected HEK 293Ts was fixed using 2% formaldehyde in phosphate buffer saline (PBS) for 30 min followed by thorough washing. Fixed cells were permeabilized at room temperature (RT) using 0.1% Triton X-100/PBS for 15 min and later blocked using 2% BSA dissolved in PBS at RT for 30 min. Proceeding to immunostaining, primary antibodies anti-RBD (Genetex, GTX135385, 1:200, USA) and anti-alphavirus (sc-293153, SantaCruz, USA, 1:200) was added to the cells and kept in 37°C humidified incubator for 1 h followed by incubation with fluorescein isothiocyanate (FITC)-conjugated goat anti-mouse secondary antibody (Invitrogen, USA,1:500 dilution) and Alexa Fluor 594 conjugated goat anti-mouse (Invitrogen, USA, 1:500 dilution) for 1 h with gentle shaking. The cells were counter stained using 4′,6-diamidino-2-phenylindole (DAPI, Sigma-Aldrich, USA) and incubated for 15 min at RT. The images were then processed using DAPI, Alexa flour 594 and FITC channels at 20X resolution using EVOS FL Auto Imaging System (Thermo scientific, USA). Expression of VLP proteins in purified VLP samples were confirmed by SDS-PAGE and RBD-E2 fusion protein was validated by western blot using anti-RBD antibody.

### 3.5. Cryo-electron microscopy and Differential Light Scattering

To examine the formation of VLPs, its size and uniformity, cryo-electron microscopy and differential light scattering (DLS) was performed. 6 μl of purified ChAC-VLP sample was adsorbed onto glow discharged quantifoil R2/2 carbon-coated 200 mesh gold grid, which was snap-frozen in liquid ethane using a Vitrobot (Mark IV), and grids were stored in liquid nitrogen for later visualization. Images were collected using a transmission electron microscope (FEI Technai G2 F20) at 200 kV at a magnification of 50,000 X. Scale bars, 100 nm. To perform DLS, 10 µL of purified ChAC-VLP was resuspended in 1 mL of TNE buffer and analysed by Malvern Zetasizer instrument. The highest intensity is observed as peak 1 with a hydrodynamic diameter of 158.9 nm.

### 3.6. Plaque reduction assay and qRT-PCR for CHIKV and SARS-CoV-2

To assess multimeric epitope display and antigenicity of RBD-E2 fusion protein on the surface of ChAC-VLPs, competitive binding is assessed by *in-vitro* experiments, i.e conventional plaque assay and qRT-PCR.

ChAC-VLP competitive plaque assay is performed on Vero cells which was maintained in DMEM supplemented with 10% heat inactivated FBS in 24-well plates. Before CHIKV infection with MOI of 1.0, Vero cells were pre-incubated with different concentrations of ChAC-VLP for 15 min to allow ChAC-VLPs to attach to the Vero cell surface. After preincubation, CHIKV dilutions was added into the wells. Following infection with CHIKV for 1 h, the infection media was removed and fresh DMEM supplemented with 2% FBS was added to the wells and plate was kept in humidified incubator at 37°C with 5% CO_2_ for 24 h. After 24 h, the virus containing supernatant is collected for later to perform plaque assay and, 200 µL triZol (TakaraBio) is added in each well for RNA isolation. For performing plaque assay, Vero cells were seeded on a 24-well plate. 10-fold serial dilutions of the viral supernatant was added to the Vero cells following infection. After 1.0 h, the infection media was removed, and cells were washed with PBS twice to remove any unbound CHIKV. Overlay media containing 1:1 ratio of 2X MEM (with 5% FBS) and carboxymethyl cellulose (CMC) (2%) was added to the wells and incubated for 48 h. Later the overlay media was removed and cells were fixed using fixing solution (10% formaldehyde). Cells were stained with 1% crystal violet and viral plaques were visualized and counted for assessment of competitive inhibition of CHIKV by ChAC-VLP. The number of plaques were counted and a decrease in viral titre was determined as PFU/mL (plaques forming unit per millilitre) and plotted on graph to determine percentage reduction in viral titre from experiments done in triplicates (Figure 2A). Similarly, quantitative RT-PCR (qRT-PCR) was performed to measure the reduction in intracellular viral mRNA of CHIKV in ChAC-VLP treated and non-treated cells. Total RNA was isolated from the 24 hpi cells CHIKV using Trizol. cDNA synthesis was performed using Accuscript High-fidelity cDNA Synthesis Kit (Agilent technologies, USA) according to manufacturer’s protocol. Actin RNA and viral RNA were measured using the KAPA SYBR fast universal qPCR kit (Sigma-Aldrich, US) on the QuantStudio real-time PCR system (Applied Biosystems, Carlsbad, CA) following manufacturer’s protocol. The results were confirmed by melting curve analysis. Experiment was performed in triplicate and ΔCt method was used to calculate relative values (Figure 2B). The relative fold change in viral RNA was calculated using ΔCt method (fold changed in viral RNA = 2^ΔCt) from the experiments done in triplicates.

qRT-PCR was performed to measure the intracellular viral RNA levels in SARS-CoV2/human/Ind/CAD1339/2020 (MOI 0.01) in ChAC-VLP treated cells at 3 different concentrations. The total RNA was isolated from the 48 h post infection VEeroE6 cell supernatant using COVISure (one step COVID19 Real Time PCR Kit by Genetix). The results were confirmed by melting curve analysis. Experiment was performed in triplicate and ΔΔCt method was used to calculate relative values. Calculated Ct values were converted to fold-reduction ChAC-VLP treated samples compared to control using the ΔCt method (fold changed in viral RNA = 2^ΔCt).

#### 3.6.2 ELISA for analysis of ChAC-VLP binding to ACE2

The study utilized 96-well microtiter plates (Greiner high binding) coated overnight at 4°C with 100 μg/mL of ACE2 protein (Sino Biological, 10108-H08H) that was diluted in PBS (pH 7.4). After washing the wells three times with 0.05 % Tween20 in PBS, the wells were blocked for 1 h at 37°C using 0.2% BSA diluted in PBS. The blocking solution was then removed and varying concentration of ChAC-VLP dissolved in 1X PBS (pH 7.4) was added to the wells and incubated for 2 h at 37°C. To serve as a negative control, only PBS was used for ACE2 coated wells. After incubation, wells were washed thrice with 0.05 % Tween20 in 1X PBS to remove any unbound VLP. 100 µL of anti-RBD antibody (1:1000) (Genetex) was added to the wells for 2 h at 37°C. The binding was detected by incubating the wells HRP tagged anti-mouse secondary antibody following addition of 100 μl of 1X TMB substrate (Genie) for 10 min, the reaction was then stopped using 1N H_2_SO_4_ and absorbance at 405 nm was measured using a microplate absorbance reader (BioTek instrument) to quantify the binding of the ChAC-VLPs to the ACE2 protein.

### 3.7. Animal Immunization and Ethics Statement

Immunogenicity of ChAC-VLP was evaluated in 6-week-old Swiss albino mice. A total of 4 mice were immunized by prime and two boost immunization through the intramuscular route. Two mice were taken as control and injected with vehicle only. For the first immunization (Prime), mice were injected by 15 µg ChAC-VLP on day 0 and in the 1st booster dose, mice were injected by 7 µg ChAC-VLP on day 14th, for the second booster dose, mice were injected with 5 µg of ChAC-VLP. Alum (Thermo scientific, USA) was used as adjuvant in the both immunizations. Mice were bled after 7 days of 3^rd^ immunization for collection of serum and later use. Animal ethics permission was obtained from ICAR-IVRI (IAEC/26.07.2021/S3).

### 3.8. Plaque Reduction Neutralization Test (PRNT) for CHIKV using ChAC-VLP immunized mice serum

Clinical CHIKV (strain 119067) (42) was propagated in Vero cells and pfu/mL was determined using conventional plaque assay method described previously (41). The assay was performed in duplicates using 24-well tissue culture plates (Genaxy, India). Serial dilutions of immunized and control mice serum samples were prepared in a 96-well plate starting from 1:10 in PBS. 40-50 plaque forming units of CHIKV was used for PRNT assay. Virus/serum interaction was performed for 1h in humidified incubator at 37°C. Proceeding to evaluation of PRNT, the virus/serum mix was added to the susceptible Vero cells and incubated for 1 h in humidified incubator at 37°C. Next, overlay media (1% carboxyl methyl cellulose with 1X MEM substituted with 5% FBS) was added to the cells and again incubated for 72 h. Plaques were visualized and counted using crystal violet solution. The percentage inhibition of virus infectivity is plotted as a function of serum dilution and PRNT50 values for the inhibition curves are calculated by GraphPad PRISM 5 Software using non-linear regression.

### 3.9. ELISA and Surface Plasmon Resonance (SPR) to detect antibody binding of post-immunization sera to recombinant RBD

High protein binding 96-well ELISA plate (Greiner bio-one, USA) was coated with 1 µg/well of RBD protein (home-made, yeast expressed) followed by blocking with 200 µL of TSTA buffer (2% bovine serum albumin (BSA), 0.9% NaCl, 0.05% Tween 20, 50 mM of Tris, pH 7.6) (49) for 2 h at 37°C. The mice serum was tenfold diluted in PBS from 1:10 to 1:1280. Diluted serum was added to the RBD-coated wells and the plate was incubated for 2 h. After washing the secondary anti-mouse HRP-labelled antibody (RD Biosystems, 1:5000) was added to the wells. Washing was performed to remove any unbound secondary antibody, followed by the addition of 100 µl/well TMB (Genei, USA). The TMB reaction was stopped by 1N H_2_SO_4_ and the absorbance was read at 450 nm.

The binding of anti-RBD antibody in post-immunization mice sera to the yeast purified RBD was analysed by SPR using a Biacore T200 system (GE Healthcare) at 25°C. Recombinant His-tagged RBD (10 µg/ml) was captured on Ni-NTA chip in 1X Phosphate buffer saline (1X PBS) running buffer. Followed by injection of 1:10 PBS diluted serum samples of non-immunized (Control) and ChAC-VLP immunized mice at a flow rate of 50 mL/min (120-s contact duration) for the association and disassociation was performed over a 600-s interval. Responses from the protein surface were corrected for the response from a mock surface.

## 4. Discussion

It is not unusual to have an acute viral infection with long-term effects. The similarities between the long-lasting manifestations of COVID-19 and those of post-chikungunya status of patients particularly in terms of decreased quality of life, and medico-social effects are surprising (50). Soon after SARS-CoV-2 emerged at the start of the twenty-first century, it was discovered that the spike (S) protein was the virus’ immunodominant antigen, particularly in its prefusion [native] configuration (51). An examination of patient antibody infected with SARS-CoV-2 revealed that the receptor-binding region of the S1 subunit is primary the target of binding and neutralizing antibodies (52). CHIKV and SARS-CoV-2 both are enveloped RNA viruses, although CHIKV infection can be fatal in very rare circumstances. The envelope glycoprotein E2 is primarily responsible for CHIKV attachment to the cell. Consequently, E2 is the immunodominant protein of CHIKV and hence the principal target of neutralizing antibodies (53, 54). There are currently no effective preventive vaccination strategies or antivirals against CHIKV infection. A need for chimeric vaccine has arisen due to global COVID-19 pandemic and unavailability of effective vaccine or antivirals for CHIKV infections. Taking into consideration the global challenge to combat any future CHIKV and SARS-CoV-2 infections, in this study, we have designed, expressed and purified, the immunogenic ChAC-VLPs, a chimeric CHIK-VLP expressing surface on its surface, fusion protein of SARS-CoV-2 RBD and CHIKV E2 glycoprotein. To the best of our knowledge, this is the first study to implement chimeric VLP technology for prevention of chikungunya virus and SARS-CoV-2.

SARS-CoV-2 Spike receptor binding domain is known to be highly glycosylated (55). The Viral envelope glycoproteins or spike proteins used in serological tests and vaccines which detect the neutralizing antibodies are often heavily glycosylated. Particularly, SARS-CoV-2 spike trimer has 66 N-glycans mainly by host-derived glycans (55). Furthermore glycosylation also partially affected binding with ACE2 which is the major receptor that interacts with RBD (55). Similarity for CHIKV, it has been reported that neutralizing antibodies detected the E2 aa 229 to 234 region in the ASR (acid-sensitive region) inhibited its conformational shift, which is required for the virus to start the process of fusion and entry into the cells (56). Studies on the n-glycosylation pattern on E1 and E2 glycoprotein has revealed high dependence on host cell expression system for proper glycosylation (57). Therefore, we chose HEK293Ts expression system for production and purification of ChAC-VLPs. The structural integrity of ChAC-VLPs was studied using cryo-electron microscopy and DLS. The results indicated that the insertions did not affect the VLP assembly. To further validate the expression of CHIKV structural proteins and RBD of SARS-CoV-2 in purified ChAC-VLPs, western blot, SDS PAGE and Indirect IFA was performed (Figure 1). The expressed E3-RBD-E2 fusion protein band from purified ChAC-VLPs migrated at ∼75 kDa, while E1 and Capsid migrated at ∼50 kDa and 35 kDa respectively. Similar band pattern was observed in western blot using anti-RBD antibody (Figure 1C). Furthermore, the efficient binding of commercially available purified ACE2 protein (Expressed in HEK293T cells) to ChAC-VLPs was demonstrated by ELISA, which was clear indicative of RBD in its correct conformation. The binding of the SARS-CoV-2 spike protein to the ACE2 receptor is a critical step in the process of entering target cells. Figure 2D, mimics the SARS-CoV-2 interaction with the ACE2 receptor of the host cell in an ELISA format as previously shown by researchers for surrogate virus neutralization test for SARS-CoV-2 [20]. To further validate the presence of immunogenic RBD-E2 epitope on the surface of ChAC-VLPs, competitive binding of ChAC-VLPs to the host cell receptors for entry of CHIKV and SARS-CoV-2, Plaque reduction assay and qRT-PCR was performed. The results of both plaque assay and qRT PCR suggest that ChAC-VLPs effectively competes with receptors for CHIKV and SARS-CoV-2 when studies *in vitro* (Figure 2). RBD in ChAC-VLP is available as antigenic epitope and competes for the host cell entry binding sites with the SARS-CoV2/human/Ind/CAD1339/2020 virus. >95% inhibition in viral titre have been observed in case of both the pathogenic virus at the highest concentration of 25.0 µg/mL of ChAC-VLPs (Figure 2C). Similarily, significant viral titer reduction is observed for CHIKV in ChAC-VLP treated cells (Figure 2A,B).

The promising results obtained from *in-vitro* experiments on CHIKV and SARS-CoV-2, prompted us to explore the immunogenicity assessment of ChAC-VLPs in mice. Four mice were immunized with ChAC-VLPs dissolved in Alum+PBS and two were treated as control (Alum+PBS). The sera were collected after 3^rd^ immunization dose (2^nd^ Booster). The prime dose amount was kept at 15 µg, following 1^st^ booster dose of 7.0 µg and 2^nd^ booster dose of 5.0 µg (Figure 3A). Expression and purification of ChAC-VLP was particularly challenging when it comes to adherent expression system and transiently transfected cells. ChAC-VLP expressing cell line in a suspension system, will improve its scalability and increased dose of ChAC-VLP might induce better immunogenic responses although potent immune response is observed at lower doses of ChAC-VLPs. Our goal was asses immunized mice sera for antibodies raised against CHIKV and RBD of SARS-CoV-2. Thus, PRNT50 was performed to detect neutralizing antibodies against CHIKV *in-vitro*. It was encouraging to observe PRNT50 of 1:25 was observed for CHIKV (Figure 3B) strongly suggesting that ChAC-VLPs have induced effective neutralizing immune response in mice for CHIKV (Clinical strain). Assessment of ChAC-VLPs to induce antibody generation against RBD was analysed by ELISA and SPR. In our study, we report that ELISA and SPR results are in coherence for serum antibodies against RBD protein. The IgG antibody (Ab) titers were calculated as the endpoint dilution that remains positively detectable for SARS-CoV-2 RBD starting at 1:40 serum antibody dilution of ChAC-VLP immunized mice (Figure 3C). ELISA is cost effective and comparatively easier to perform compared to SPR. Although, SPR evaluates real time binding of antibodies to the immobilised antigen, total binding of antibodies to antigen is expressed in the form of response unites (max RU), along with very low usage of sera samples. The purified RBD protein was captured on Ni-NTA chip in its native form up to RU value of ∼ 250. Immobilisation in the native form of protein is an advantage over other techniques such as ELISA. In ELISA proteins are coated in the plate which can cause partial denaturation of antigens. When PBS diluted sera was injected on immobilised RBD, the response unit for 2 of the immunized mice (Immunized 2 and 4) was twofold higher as compared to control, whereas the other two mice (Immunised 1 and 3) showed threefold higher reading compared to the control sera. Prominent difference in the reactivity towards RBD of immunized and non-immunised mice sera was observed in the form of response units. Thus, the SPR results indicate immunization has induced anti-RBD antibodies in the sera of mice. (Figure 4)

## Conclusion

In conclusion, the development of a chimeric chikungunya virus-like particles vaccine that displays the RBD of SARS-CoV-2 and the envelop (E2) glycoprotein of CHIKV is a promising strategy for bivalent vaccination against both the viruses. We have utilized an alphavirus structural proteins backbone to successfully integrate and display on the surface of VLPs, fusion protein of RBD-E2 as these being the immunodominant proteins of SARS-CoV-2 and CHIKV respectively. Our experiments have shown successful generation of ChAC-VLPs which have the ability to elicit significant neutralizing activity against both CHIKV and SARS-CoV-2 in mice. This work highlights the potential of ChAC-VLPs as a hybrid CHIKV and SARS-CoV-2 vaccine platform that may be a valuable addition to the worldwide vaccine development and navigation in enveloped virus-like particle (eVLP) domain.

## Acknowledgements

ST thank Department of Biotechnology, Govt of India for supporting Bioinformatics Center at IIT Roorkee (reference number BT/PR40141/BTIS/137/16/2021). V.A.S. and S.N acknowledges the Ministry of Human Resource Development, (MHRD), for financial support.

## References

1. Tariq H, Batool S, Asif S, Ali M, Abbasi BH. 2022. Virus-Like Particles: Revolutionary Platforms for Developing Vaccines Against Emerging Infectious Diseases. Front Microbiol https://doi.org/10.3389/fmicb.2021.790121.

2. Qian C, Liu X, Xu Q, Wang Z, Chen J, Li T, Zheng Q, Yu H, Gu Y, Li S, Xia N. 2020. Recent Progress on the Versatility of Virus-Like Particles. Vaccines 2020, Vol 8, Page 139 8:139.

3. Zhang P, Narayanan E, Liu Q, Tsybovsky Y, Boswell K, Ding S, Hu Z, Follmann D, Lin Y, Miao H, Schmeisser H, Rogers D, Falcone S, Elbashir SM, Presnyak V, Bahl K, Prabhakaran M, Chen X, Sarfo EK, Ambrozak DR, Gautam R, Martin MA, Swerczek J, Herbert R, Weiss D, Misamore J, Ciaramella G, Himansu S, Stewart-Jones G, McDermott A, Koup RA, Mascola JR, Finzi A, Carfi A, Fauci AS, Lusso P. 2021. A multiclade env–gag VLP mRNA vaccine elicits tier-2 HIV-1-neutralizing antibodies and reduces the risk of heterologous SHIV infection in macaques. Nat Med 2021 2712 27:2234–2245.

4. Mazumder S, Rastogi R, Undale A, Arora K, Arora NM, Pratim B, Kumar D, Joseph A, Mali B, Arya VB, Kalyanaraman S, Mukherjee A, Gupta A, Potdar S, Roy SS, Parashar D, Paliwal J, Singh SK, Naqvi A, Srivastava A, Singh MK, Kumar D, Bansal S, Rautray S, Saini M, Jain K, Gupta R, Kundu PK. 2021. PRAK-03202: A triple antigen virus-like particle vaccine candidate against SARS CoV-2. Heliyon 7:e08124.

5. Lu J, Lu G, Tan S, Xia J, Xiong H, Yu X, Qi Q, Yu X, Li L, Yu H, Xia N, Zhang T, Xu Y, Lin J. 2020. A COVID-19 mRNA vaccine encoding SARS-CoV-2 virus-like particles induces a strong antiviral-like immune response in mice. Cell Res https://doi.org/10.1038/s41422-020-00392-7.

6. Xu R, Shi M, Li J, Song P, Li N. 2020. Construction of SARS-CoV-2 Virus-Like Particles by Mammalian Expression System. Front Bioeng Biotechnol 8:862.

7. Singh K, Marasini B, Chen X, Ding L, Wang J-J, Xiao P, Villinger F, Spearman P. 2020. A Bivalent, Spherical Virus-Like Particle Vaccine Enhances Breadth of Immune Responses against Pathogenic Ebola Viruses in Rhesus Macaques. J Virol 94.

8. Warfield KL, Bosio CM, Welcher BC, Deal EM, Mohamadzadeh M, Schmaljohn A, Aman MJ, Bavari S. 2003. Ebola virus-like particles protect from lethal Ebola virus infection. Proc Natl Acad Sci U S A 100:15889–15894.

9. Boigard H, Cimica V, Galarza JM. 2018. Dengue-2 virus-like particle (VLP) based vaccine elicits the highest titers of neutralizing antibodies when produced at reduced temperature. Vaccine 36:7728–7736.

10. Liu Y, Zhou J, Yu Z, Fang D, Fu C, Zhu X, He Z, Yan H, Jiang L. 2014. Tetravalent recombinant dengue virus-like particles as potential vaccine candidates: Immunological properties. BMC Microbiol 14.

11. Urakami A, Ngwe Tun MM, Moi ML, Sakurai A, Ishikawa M, Kuno S, Ueno R, Morita K, Akahata W. 2017. An Envelope-Modified Tetravalent Dengue Virus-Like-Particle Vaccine Has Implications for Flavivirus Vaccine Design. J Virol 91.

12. Bennett SR, McCarty JM, Ramanathan R, Mendy J, Richardson JS, Smith J, Alexander J, Ledgerwood JE, de Lame PA, Royalty Tredo S, Warfield KL, Bedell L. 2022. Safety and immunogenicity of PXVX0317, an aluminium hydroxide-adjuvanted chikungunya virus-like particle vaccine: a randomised, double-blind, parallel-group, phase 2 trial. Lancet Infect Dis 22:1343–1355.

13. Akahata W, Yang ZY, Andersen H, Sun S, Holdaway HA, Kong WP, Lewis MG, Higgs S, Rossmann MG, Rao S, Nabel GJ. 2010. A virus-like particle vaccine for epidemic Chikungunya virus protects nonhuman primates against infection. Nat Med 16:334–338.

14. M RB, ell, Flick R. 2011. Virus-Like Particle-Based vaccines for Rift Valley Fever Virus. J Bioterrorism Biodefense 2012 00 s1:1–5.

15. Christiansen D, Earnest-Silveira L, Grubor-Bauk B, Wijesundara DK, Boo I, Ramsland PA, Vincan E, Drummer HE, Gowans EJ, Torresi J. 2019. Pre-clinical evaluation of a quadrivalent HCV VLP vaccine in pigs following microneedle delivery. Sci Reports 2019 91 9:1–13.

16. Torresi J. 2017. The Rationale for a Preventative HCV Virus-Like Particle (VLP) Vaccine. Front Microbiol 8:2163.

17. Kushnir N, Streatfield SJ, Yusibov V. 2012. Virus-like particles as a highly efficient vaccine platform: Diversity of targets and production systems and advances in clinical development. Vaccine 31:58–83.

18. Crisci E, Bárcena J, Montoya M. 2012. Virus-like particles: The new frontier of vaccines for animal viral infections. Vet Immunol Immunopathol 148:211–225.

19. Zepeda-Cervantes J, Ramírez-Jarquín JO, Vaca L. 2020. Interaction Between Virus-Like Particles (VLPs) and Pattern Recognition Receptors (PRRs) From Dendritic Cells (DCs): Toward Better Engineering of VLPs. Front Immunol 11:1100.

20. Nooraei S, Bahrulolum H, Hoseini ZS, Katalani C, Hajizade A, Easton AJ, Ahmadian G. 2021. Virus-like particles: preparation, immunogenicity and their roles as nanovaccines and drug nanocarriers. J Nanobiotechnology 2021 191 19:1–27.

21. Plotkin S. 2014. History of vaccination. Proc Natl Acad Sci 111:12283–12287.

22. Brisse M, Vrba SM, Kirk N, Liang Y, Ly H. Emerging Concepts and Technologies in Vaccine Development https://doi.org/10.3389/fimmu.2020.583077.

23. Dai S, Wang H, Deng F. 2018. Advances and challenges in enveloped virus-like particle (VLP)-based vaccines. J Immunol Sci 2:36–41.

24. Lua LHL, Connors NK, Sainsbury F, Chuan YP, Wibowo N, Middelberg APJ. 2014. Bioengineering virus-like particles as vaccines. Biotechnol Bioeng 111:425–440.

25. Lin S-Y, Chung Y-C, Chiu H-Y, Chi W-K, Chiang B-L, Hu Y-C. 2014. Evaluation of the stability of enterovirus 71 virus-like particle. J Biosci Bioeng 117:366–371.

26. Donaldson B, Lateef Z, Walker GF, Young SL, Ward VK. 2018. Virus-like particle vaccines: immunology and formulation for clinical translation https://doi.org/10.1080/14760584.2018.1516552.

27. Mejía-Méndez JL, Vazquez-Duhalt R, Hernández LR, Sánchez-Arreola E, Bach H. 2022. Virus-like Particles: Fundamentals and Biomedical Applications. Int J Mol Sci 23:8579.

28. Lei X, Cai X, Yang Y. 2020. Genetic engineering strategies for construction of multivalent chimeric VLPs vaccines. https://doi.org/101080/1476058420201738227 19:235–246.

29. Reisinger EC, Tschismarov R, Beubler E, Wiedermann U, Firbas C, Loebermann M, Pfeiffer A, Muellner M, Tauber E, Ramsauer K. 2019. Immunogenicity, safety, and tolerability of the measles-vectored chikungunya virus vaccine MV-CHIK: a double-blind, randomised, placebo-controlled and active-controlled phase 2 trial. Lancet (London, England) 392:2718–2727.

30. Boxus M, Fochesato M, Miseur A, Mertens E, Dendouga N, Brendle S, Balogh KK, Christensen ND, Giannini SL. 2016. Broad Cross-Protection Is Induced in Preclinical Models by a Human Papillomavirus Vaccine Composed of L1/L2 Chimeric Virus-Like Particles. J Virol 90:6314–6325.

31. B H, C S, S S-K, C J, N C, R K. 2017. Chimeric L2-Based Virus-Like Particle (VLP) Vaccines Targeting Cutaneous Human Papillomaviruses (HPV). PLoS One 12.

32. Phelps DK, Speelman B, Post CB. 2000. Theoretical studies of viral capsid proteins. Curr Opin Struct Biol 10:170–173.

33. Sasagawa T, Pushko P, Steers G, Gschmeissner SE, Nasser Hajibagheri MA, Finch J, Crawford L, Tommasino M. 1995. Synthesis and assembly of virus-like particles of human papillomaviruses type 6and Type 16 in fission yeast Schizosaccharomyces pombe. Virology 206:126–135.

34. Haynes JR. 2014. Influenza virus-like particle vaccines. https://doi.org/101586/erv098 8:435–445.

35. Chen BJ, Leser GP, Morita E, Lamb RA. 2007. Influenza Virus Hemagglutinin and Neuraminidase, but Not the Matrix Protein, Are Required for Assembly and Budding of Plasmid-Derived Virus-Like Particles. J Virol 81:7111–7123.

36. Fatma B, Kumar R, Singh VA, Nehul S, Sharma R, Kesari P, Kuhn RJ, Tomar S. 2020. Alphavirus capsid protease inhibitors as potential antiviral agents for Chikungunya infection. Antiviral Res 179:104808.

37. Pialoux G, Gaüzère BA, Jauréguiberry S, Strobel M. 2007. Chikungunya, an epidemic arbovirosis. Lancet Infect Dis 7:319–327.

38. Yap ML, Klose T, Urakami A, Hasan SS, Akahata W, Rossmann MG. 2017. Structural studies of Chikungunya virus maturation. Proc Natl Acad Sci 114:13703–13707.

39. Simon F, Watson H, Meynard J-B, Santi VP de, Tournier J-N. 2021. What chikungunya teaches us about COVID-19. Lancet Infect Dis 21:1070–1071.

40. Singh VA, Kumar CS, Khare B, Kuhn RJ, Banerjee M, Tomar S. 2023. Surface decorated reporter-tagged chikungunya virus-like particles for clinical diagnostics and identification of virus entry inhibitors. Virology 578:92–102.

41. Choudhary S, Neetu N, Singh VA, Kumar P, Chaudhary M, Tomar S. 2021. Chikungunya virus titration, detection and diagnosis using N-Acetylglucosamine (GlcNAc) specific lectin based virus capture assay. Virus Res 302:198493.

42. Singh H, Mudgal R, Narwal M, Kaur R, Singh VA, Malik A, Chaudhary M, Tomar S. 2018. Chikungunya virus inhibition by peptidomimetic inhibitors targeting virus-specific cysteine protease. Biochimie 149:51–61.

43. Fernandes Q, Inchakalody VP, Merhi M, Mestiri S, Taib N, Moustafa Abo El-Ella D, Bedhiafi T, Raza A, Al-Zaidan L, Mohsen MO, Yousuf Al-Nesf MA, Hssain AA, Yassine HM, Bachmann MF, Uddin S, Dermime S. 2022. Emerging COVID-19 variants and their impact on SARS-CoV-2 diagnosis, therapeutics and vaccines. Ann Med 54:524–540.

44. Khan WH, Hashmi Z, Goel A, Ahmad R, Gupta K, Khan N, Alam I, Ahmed F, Ansari MA. 2021. COVID-19 Pandemic and Vaccines Update on Challenges and Resolutions. Front Cell Infect Microbiol 11.

45. Greaney AJ, Starr TN, Barnes CO, Weisblum Y, Schmidt F, Caskey M, Gaebler C, Cho A, Agudelo M, Finkin S, Wang Z, Poston D, Muecksch F, Hatziioannou T, Bieniasz PD, Robbiani DF, Nussenzweig MC, Bjorkman PJ, Bloom JD. 2021. Mapping mutations to the SARS-CoV-2 RBD that escape binding by different classes of antibodies. Nat Commun 2021 121 12:1–14.

46. Jin J, Sherman MB, Chafets D, Dinglasan N, Lu K, Lee TH, Carlson LA, Muench MO, Simmons G. 2018. An attenuated replication-competent chikungunya virus with a fluorescently tagged envelope. PLoS Negl Trop Dis 12:e0006693.

47. Tan CW, Chia WN, Qin X, Liu P, Chen MIC, Tiu C, Hu Z, Chen VCW, Young BE, Sia WR, Tan YJ, Foo R, Yi Y, Lye DC, Anderson DE, Wang LF. 2020. A SARS-CoV-2 surrogate virus neutralization test based on antibody-mediated blockage of ACE2–spike protein–protein interaction. Nat Biotechnol 2020 389 38:1073–1078.

48. Jose J, Tang J, Taylor AB, Baker TS, Kuhn RJ. 2015. Fluorescent Protein-Tagged Sindbis Virus E2 Glycoprotein Allows Single Particle Analysis of Virus Budding from Live Cells. Viruses 7:6182.

49. David A Ben, Diamant E, Dor E, Barnea A, Natan N, Levin L, Chapman S, Mimran LC, Epstein E, Zichel R, Torgeman A. 2021. Identification of SARS-CoV-2 Receptor Binding Inhibitors by In Vitro Screening of Drug Libraries. Molecules 26.

50. Simon F, Watson H, Meynard JB, de Santi VP, Tournier JN. 2021. What chikungunya teaches us about COVID-19. Lancet Infect Dis 21:1070–1071.

51. Harvey WT, Carabelli AM, Jackson B, Gupta RK, Thomson EC, Harrison EM, Ludden C, Reeve R, Rambaut A, Peacock SJ, Robertson DL. 2021. SARS-CoV-2 variants, spike mutations and immune escape. Nat Rev Microbiol 2021 197 19:409–424.

52. Premkumar L, Segovia-Chumbez B, Jadi R, Martinez DR, Raut R, Markmann AJ, Cornaby C, Bartelt L, Weiss S, Park Y, Edwards CE, Weimer E, Scherer EM, Rouphael N, Edupuganti S, Weiskopf D, Tse L V., Hou YJ, Margolis D, Sette A, Collins MH, Schmitz J, Baric RS, de Silva AM. 2020. The receptor-binding domain of the viral spike protein is an immunodominant and highly specific target of antibodies in SARS-CoV-2 patients. Sci Immunol 5:8413.

53. Burt FJ, Rolph MS, Rulli NE, Mahalingam S, Heise MT. 2012. Chikungunya: A re-emerging virus. Lancet 379:662–671.

54. Tumkosit U, Siripanyaphinyo U, Takeda N, Tsuji M, Maeda Y, Ruchusatsawat K, Shioda T, Mizushima H, Chetanachan P, Wongjaroen P, Matsuura Y, Tatsumi M, Tanaka A. 2020. Anti-Chikungunya Virus Monoclonal Antibody That Inhibits Viral Fusion and Release. J Virol 94:252–272.

55. Watanabe Y, Allen JD, Wrapp D, McLellan JS, Crispin M. 2020. Site-specific glycan analysis of the SARS-CoV-2 spike. Science 369:330.

56. Akahata W, Nabel GJ. 2012. A Specific Domain of the Chikungunya Virus E2 Protein Regulates Particle Formation in Human Cells: Implications for Alphavirus Vaccine Design. J Virol 86:8879–8883.

57. Lancaster C, Pristatsky P, Hoang VM, Casimiro DR, Schwartz RM, Rustandi R, Ha S. 2016. Characterization of N-glycosylation profiles from mammalian and insect cell derived chikungunya VLP. J Chromatogr B Analyt Technol Biomed Life Sci 1032:218–223.

